# Integrative analysis of relative abundance data and presence-absence data of the microbiome using the LDM

**DOI:** 10.1101/2022.01.14.476390

**Authors:** Zhengyi Zhu, Glen A. Satten, Yi-Juan Hu

## Abstract

**Summary:** We previously developed LDM for testing hypotheses about the microbiome that performs the test at both the community level and the individual taxon level. LDM can be applied to relative abundance data and presence-absence data separately, which work well when associated taxa are abundant and rare, respectively. Here we propose an omnibus test based on LDM that allows simultaneous consideration of data at different scales, thus offering optimal power across scenarios with different association mechanisms. The omnibus test is available for the wide range of data types and analyses that are supported by LDM.

**Availability and Implementation:** The omnibus test has been added to the R package LDM, which is available on GitHub at https://github.com/yijuanhu/LDM.

**Supplementary information:** Supplementary data are available at Bioinformatics online.

## Introduction

In recent years, there have been significant advances in statistical methodologies for analyzing associations of the microbiome with traits of interest such as clinical outcomes and environmental factors. It is generally accepted that the analysis based on relative abundance data will work best when associated taxa are abundant, while the analysis based on presenceabsence data will work best when associated taxa are less abundant. Because the association mechanism is not known a priori, one strategy is to conduct analysis at each taxon scale separately and combine the results in an omnibus test. For example, PERMANOVA-S [1] and MiRKAT-O [2] provided omnibus tests that combine results from analyzing multiple distance matrices, such as the weighted and unweighted UniFrac distances in a phylogenetic-tree-based approach or Bray-Curtis and Jaccard distances in a non-tree-based approach; note that the weighted UniFrac and Bray-Curtis distances are based on relative abundance data while the unweighted UniFrac and Jaccard distances are based on presence-absence data.

We previously developed LDM [3–7] for testing hypotheses about the microbiome, which not only performs the test at the community level, like PERMANOVA [8] and MiRKAT [2], but also at the individual taxon level with false discovery rate (FDR) control. LDM is built upon a linear model that regresses the taxon data on covariates including the confounding covariates that we wish to adjust for and the traits that we wish to test. The inference is based on permutation. These features made LDM highly versatile, applicable to a wide variety of data types and analyses that are listed in Table 1.

**Table 1.** Data types and analyses that LDM supports

| Reference | Taxon scale | Trait type | Sample structure | Hypothesis |
| --- | --- | --- | --- | --- |
| Hu and Satten (2020) [3] | relative abundance,<br>arcsin-root transformed | continuous,<br>discrete,<br>multivariate | independent,<br>correlated | association,<br>interaction |
| Hu et al. (2021) [4] | presence-absence |  |  |  |
| Hu et al. (2022) [7] |  | time-to-event |  |  |
| Zhu et al. (2021) [5] |  |  | matched sets |  |
| Yue and Hu (2021) [6] |  |  |  | mediation |

Note: The proposed omnibus test is available for all supported analyses.

LDM was initially developed for taxon data at the relative abundance scale and the arcsin-root-transformed relative abundance scale (which is variance-stabilizing for Multinomial and Dirichlet-Multinomial count data), and also offered an omnibus test that combined the results of the two taxon scales [3]. We have shown that LDM applied to the untransformed data worked better when associated taxa were abundant and LDM applied to the transformed data worked better when associated taxa were less abundant. More recently, we made an extension of LDM for analyzing data at the presence-absence scale [4], which accounted for variability of library size by a rarefaction-based yet non-stochastic approach that evaluated the expected LDM test statistic over all rarefaction replicates. We found that the presence-absence analysis performed better than the relative-abundance-based analysis when associated taxa were more rare. These results motivated us to develop a *new* omnibus test for LDM that combines results from all three taxon scales. Here, we present such an omnibus test at both the community level and the individual taxon level.

## Methods

### Taxon-level omnibus test

It is straightforward to construct an LDM omnibus test for each taxon. LDM-omni, the omnibus test in [3], used the minimum of the *p*-values obtained from analyzing the frequency (i.e., relative abundance) data and the arcsin-root-transformed data at each taxon as the final test statistic, and used the corresponding minima from the permutation replicates to simulate the null distribution. Now we expand the test statistic to include the *p*-value from the presence-absence analysis in the calculation of the minimum at each taxon. As in [3], we apply Sandve’s [9] sequential Monte-Carlo multiple-testing procedure to make discoveries with FDR control. We refer to the new omnibus test as LDM-omni3 with “3” indicates the “three” taxon scales.

Both LDM-omni and LDM-omni3 combine different scales of data at the *p*-value level. A completely different way to combine data would be to combine two separate lists of discoveries, each preserving FDR at some level, so that the combined list of discoveries controls the overall FDR. Kim et al. [10] gave such a method; thus, we will compare LDM-omni3 to Kim et al.’s method that combines the discovery lists of LDM-omni and the presence-absence analysis, each using half of the overall nominal FDR. We denote this test by LDM-omni-Kim.

### Community-level omnibus test

A community-level (global) version of LDM-omni3 could easily be constructed in the same way that LDM-omni combined information across the frequency and arcsin-root scales [3], by calculating an *overall F*-statistic (and corresponding *p*-value) for each scale (frequency, arcsin-root and, new for LDM-omni3, presence-absence) and choosing the scale with the minimum *p*-value. However, we also want to construct a global test that allows using the best scale *at each taxon*. Thus, we consider various *p*-value combination methods to combine the taxon-level LDM-omni3 *p*-values into a statistic we could *add* to the global LDM-omni3 test; we initially considered the minimum *p*-value over taxa, as well as the Cauchy [11], Harmonic-mean (HM) [12], Fisher’s, and Stouffer’s methods.

An immediate problem in combining permutation *p*-values calculated using *B* replicate datasets is that the *p*-values have the minimum achievable value 1/(*B* + 1) [13] and thus cannot well estimate the tail probability of the test statistic [14]. This can greatly diminish the power of most *p*-value combination methods, which highly depend on the smallest *p*-values. To overcome this, we propose to apply the combination method to the “analytical” *p*-value, which is the tail probability of the *F*-statistic for each taxon compared to the corresponding *F* distribution. While the *F* statistics for many taxa follow the *F* distribution, others tend to have smaller empirical variances, in which case we scale the *F* statistics to have the expected variance of the *F* distribution. We then calculate the taxon-level “analytic” *p*-value for each scale of data, take the minimum of the *p*-values corresponding to the three scales, and combine these minimum “analytic” *p*-values across taxa using a *p*-value combination method. It is important to note here that the “analytic” *p*-values are not used as *real p*-values but rather a transformation of the taxon-level *F*-statistic to an appropriate scale for combining. The resulting statistic of a p-value combination method is assessed for significance using the permutation replicates.

For our new global LDM-omni3 test, we choose to add global tests that are based on the HM and Fisher’s *p*-value combination methods; motivation for this choice can be found in Text S1. Thus, the global LDM-omni3 test statistic is calculated as

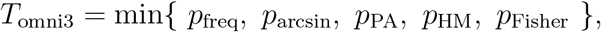

where *p*_freq_, *p*_arcsin_, and *p*_PA_ are the permutation *p*-values of the global *F*-statistics at the frequency, arcsin-root and presence-absence scales as described in [3, 4], while *p_HM_* and *p*_Fisher_ are the permutation p-values of the HM and Fisher’s combinations of the “analytic” *p*-values. The significance of *T*_omni3_ is determined by permutation. We note that, by adding *p*-value combination tests to LDM-omni3, we increase the consistency of the taxon-level and global tests, because the taxon-level LDM-omni test in [3] reports results at the scale having the smallest *p*-value for that taxon, while the global LDM-omni test in [3], like *p*_freq_, *p*_arcsin_, and *p*_PA_, is calculated using the same scale at every taxon. Finally, note that any requirement on independence of p-values when computing *p*-value combination tests is irrelevant here as inference is based on permutation.

## Results

### Simulation studies

Our simulations were based on the Dirichlet-Multinomial model and data on 856 taxa of the upper-respiratory-tract (URT) microbiome [15], both of which were also used by the LDM paper [3]. We selected five different sets of taxa to be associated with a binary or continuous trait. Ordering taxa by decreasing relative abundance, these sets are: (S1) taxa 1-10, (S2) taxa 11–50, (S3) taxa 51–200, (S4) taxa 3–5 and 11–50, and (S5) taxa 3–5 and 51–200. By design, the taxa in S1, S2, and S3 were abundant, less abundant, and relatively rare, respectively, and those in S4 and S5 were mixtures of these taxa. For each set of taxa, we considered two models for generating associations with the trait. Briefly, in Model 1, we assumed a binary trait and used different frequencies at associated taxa to simulate read count data for samples with different trait levels; in Model 2, we assumed a continuous trait and related it to associated taxa through a weighted sum of their frequencies. More detail is provided in Text S2. While Model 1 tended to create strong associations at a few taxa, Model 2 tended to simulate weak associations for all associated taxa (although the abundant taxa generally had a higher impact).

For testing individual taxa, we compared LDM-omni3 to LDM-omni and LDM-omni-Kim at the nominal FDR of 10%. To gain more insights into the relative performance of the three taxon scales, we considered the results of LDM applied to each single scale of data and call them LDM-freq, LDM-arcsin, and LDM-PA. For testing the global association, we again compared LDM-omni3 to LDM-omni. Because LDM-omni3 combined results of the five global tests, we also considered their results separately and call them LDM-freq, LDM-arcsin, LDM-PA, LDM-HM, and LDM-Fisher.

Figure 1 shows the results of sensitivity of taxon-specific tests and power of the global test for all methods across all ten scenarios. Note that all methods controlled the FDR or type I error (Figure S1). For testing individual taxa, LDM-freq, LDM-arcsin, and LDM-PA each performed well in some scenarios but poorly in others, whereas LDM-omni3 achieved good sensitivity across all scenarios and often tracked the best-performing scale. LDM-omni3 outperformed LDM-omni-Kim in all scenarios. LDM-omni3 yielded higher and sometimes much higher sensitivity over LDM-omni when the presence-absence analysis worked well (e.g., S2-S3 in Model 1) and lost only a small amount of sensitivity otherwise. The results at the global level showed a very similar pattern. The five tests that were components of the global LDM-omni3 test each performed well in some scenarios but poorly in others, whereas the global LDM-omni3 test achieved good power across all scenarios. When LDM-omni3 gained power over LDM-omni, the power gain can be as large as 200% (e.g., S3 and S5 in Model 1); when LDM-omni3 lost power to LDM-omni, the lost was usually small.

**Figure 1:**
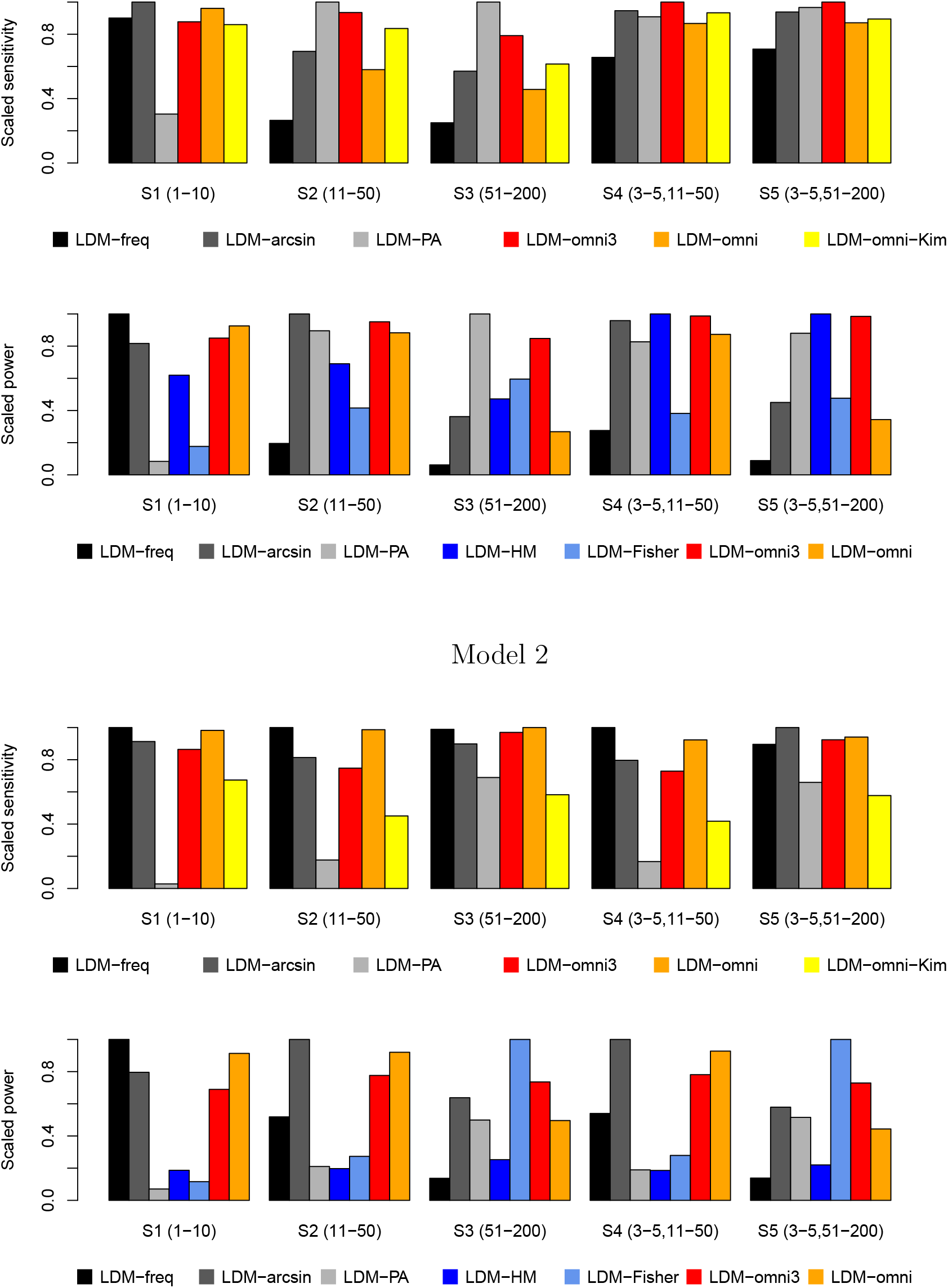
The scaled sensitivity and scaled power values were ratios against the largest value of all methods in each scenario. All results were based on 1000 replicates of data.

### Testing Association in the URT microbiome dataset

We tested the association of the URT microbiome with smoking status in the data of Charlson et al. [15], controlling for potential confounders gender and antibiotic use. More details on the dataset and the pre-processing procedures used can be found in [3]. For testing the global association, LDM-freq, LDM-arcsin, LDM-PA, LDM-HM, and LDM-Fisher yielded *p*-values 0.0077, 0.00060, 0.0075, 0.020, and 0.011, respectively. LDM-omni3 yielded the second smallest *p*-value 0.0018, which tracked the best-performing LDM-arcsin test. In this case, when LDM-freq and LDM-arcsin worked exceptionally well, it was not surprising that LDM-omni generated the small *p*-value of 0.00090.

At the OTU level, LDM-freq, LDM-arcsin, and LDM-PA detected 4, 14, and 3 OTUs, respectively, that had significant associations with smoking at nominal FDR 10%. LDM-omni3 yielded 7 detections, which included two novel OTUs compared to the 5 detections by LDM-omni. One novel OTU (OTU 411) was only detected by LDM-PA. This OTU was present in 8 smokers only and absent in all others (*p* = 0.0012 by Fisher’s exact test), confirming a strong association of the presence-absence data with smoking status. In contrast, the relative abundance data did not show much difference except for two smokers with fairly high values, although the difference was more pronounced on the arcsin-root scale (Figure S2). The other novel OTU, OTU 3954, was detected by both LDM-PA and LDM-arcsin but the latter *q*-value was just at the boundary of the FDR threshold. This OTU was absent in 6 smokers (*p* = 0.0075 by Fisher’s), also confirming a strong association at the presence-absence scale.

## References

1. Tang ZZ, Chen G, Alekseyenko AV. PERMANOVA-S: association test for microbial community composition that accommodates confounders and multiple distances. Bioinformatics. 2016;32(17):2618–2625.

2. Zhao N, Chen J, Carroll IM, Ringel-Kulka T, Epstein MP, Zhou H, et al. Testing in microbiome-profiling studies with MiRKAT, the microbiome regression-based kernel association test. The American Journal of Human Genetics. 2015;96(5):797–807.

3. Hu YJ, Satten GA. Testing hypotheses about the microbiome using the linear decomposition model (LDM). Bioinformatics. 2020;p. bbtaa260, https://doi.org/10.1093/bioinformatics/btaa260.

4. Hu YJ, Lane A, Satten GA. A rarefaction-based extension of the LDM for testing presence-absence associations in the microbiome. Bioinformatics. 2021;p. https://doi.org/10.1093/bioinformatics/btab012.

5. Zhu Z, Satten GA, Mitchell C, Hu YJ. Constraining PERMANOVA and LDM to within-set comparisons by projection improves the efficiency of analyses of matched sets of microbiome data. Microbiome. 2021;9(1):1–19.

6. Yue Y, Hu YJ. A New Approach to Testing Mediation of the Microbiome using the LDM. bioRxiv. 2021;.

7. Hu Y, Satten GA, Hu YJ. Testing associations of the microbiome with censored survival outcomes using the LDM and PERMANOVA. bioRxiv. 2021;.

8. McArdle BH, Anderson MJ. Fitting multivariate models to community data: a comment on distance-based redundancy analysis. Ecology. 2001;82(1):290—297.

9. Sandve GK, Ferkingstad E, Nygård S. Sequential Monte Carlo multiple testing. Bioinformatics. 2011;27(23):3235—3241.

10. Kim Y, Lim J, Lee JS, Jeong J. Controlling two-dimensional false discovery rates by combining two univariate multiple testing results with an application to mass spectral data. Chemometrics and Intelligent Laboratory Systems. 2018;182:149—157.

11. Liu Y, Xie J. Cauchy combination test: a powerful test with analytic p-value calculation under arbitrary dependency structures. Journal of the American Statistical Association. 2020;115(529):393—402.

12. Wilson DJ. The harmonic mean p-value for combining dependent tests. Proceedings of the National Academy of Sciences. 2019;116(4):1195—1200.

13. Besag J, Clifford P. Sequential Monte Carlo p-values. Biometrika. 1991;78(2):301—304.

14. Phipson B, Smyth GK. Permutation P-values should never be zero: calculating exact P-values when permutations are randomly drawn. Statistical applications in genetics and molecular biology. 2010;9(1).

15. Charlson ES, Chen J, Custers-Allen R, Bittinger K, Li H, Sinha R, et al. Disordered microbial communities in the upper respiratory tract of cigarette smokers. PloS one. 2010;5(12):e15216. PMCID: PMC3004851.

