## Supplementary material for "Integrative analysis of relative abundance data and presence-absence data of the microbiome using the LDM": Supplmental file

### Supplementary Materials

#### Text S1: Choosing among $p$-value combination methods

A new feature of LDM-omni3, compared to LDM-omni given in [1], is the use of statistics that aggregate the taxon-level omnibus $p$-values using a $p$-value combination method. The most commonly used $p$-value combination methods, namely, the minimum $p$-value, Cauchy [2], Harmonic-mean (HM) [3], Fisher's, and Stouffer's methods, have a decreasing emphasis on the smallest $p$-values and an increasing focus on the proportion of modest to weak signals [4, 5] in the order given here. Thus, different methods suit different scenarios; more detail is provided in Table S1.

Since each combination method has its own strength, it is desirable to further combine their results to form our new, omnibus global test. However, in our simulations, we found that: the Cauchy method always gave almost identical results as the HM method; the minimum $p$-value method performed uniformly worse than HM; and Stouffer's method performed uniformly worse than Fisher's. Stouffer's method would work better than Fisher's method when "all nulls are equally false" [4], but this is unrealistic in the microbiome setting. As a results, for our new global LDM-omni3 test, we chose to add global tests that are based on the HM and Fisher's $p$-value combination methods.

Table S1. $P$-value combination methods

| $P$-value combination method | Test statistic | Suitable scenario |
|---|---|---|
| Minimum $p$-value | $T_{\text{minP}} = \min_{j=1,\ldots,J}\{p_j\}$ | One strongest signal |
| Cauchy | $T_{\text{Cauchy}} = \sum_{j=1}^{J} \tan\{(0.5 - p_j)\pi\}$ | A very few strong signals |
| Harmonic-mean (HM) | $T_{\text{HM}} = \sum_{j=1}^{J} p_j^{-1}$ | A very few strong signals |
| Fisher's | $T_{\text{Fisher}} = -2\sum_{j=1}^{J} \log(p_j)$ | A few strong to moderate signals |
| Stouffer's | $T_{\text{Stouffer}} = \sum_{j=1}^{J} \Phi^{-1}(1 - p_j/2)$ | A large proportion of weak signals |

Note: $\{p_1, p_2, \ldots, p_J\}$ are individual $p$-values. $T_{\text{HM}}$ here is the inverse of the usual HM statistic so that $T_{\text{HM}}$ has the property of a usual test statistic that a large value corresponds to a stronger evidence against the null hypothesis. $\Phi(\cdot)$ is the standard normal cumulative distribution function.

#### Text S2: Two models for simulating microbiome-trait associations

Model 1 with a binary trait was previously considered in the LDM paper [1]. We let $X_i$ denote the trait of sample $i$ and assumed 50 samples with $X_i = 1$ and 50 with $X_i = 0$. We let $\pi_0$ be the vector of taxon frequencies estimated from the upper-respiratory-tract microbiome data; we assign $\pi_0$ to samples for which $X_i = 0$. We derived a second set of taxon frequencies $\pi_1$ by first setting $\pi_1 = \pi_0$ and then randomly permuting the frequencies in $\pi_1$ that belonged to the selected set of taxa associated with the trait, which ensured the same frequencies in $\pi_0$ and $\pi_1$ for taxa not selected. We then defined a sample-specific frequency vector as $\widetilde{\pi}(X_i|\beta) = (1 - \beta X_i)\pi_0 + \beta X_i \pi_1$, where $\beta$ can be interpreted as the effect size of the trait on the overall community composition. The strengths and directions of the effects of the trait on individual taxa were heterogeneous because the resulting frequencies at each taxon were characterized not only by $\beta$ but also by the differences between $\pi_0$ and $\pi_1$, which varied in magnitude and sign at different taxa. Because the frequency distribution was highly skewed towards zero, this model tended to create strong associations at only a very few taxa, leaving the majority weakly associated. Finally, we generated the taxon count data for each sample using the Dirichlet-Multinomial model with mean $\widetilde{\pi}(X_i|\beta)$, overdispersion 0.02, and library size sampled from $N(10000, (10000/3)^2)$ and left-truncated at 500.

Model 2 with a continuous trait is similar to the one considered in the MiRKAT paper [6]. We first generated the taxon count data for 100 samples using the same Dirichlet-Multinomial model as above except for the mean, which was set to $\pi_0$ here. Define $C_i = \sum_{j \in \mathcal{A}} \delta_j Y_{ij}/\overline{Y}_j$, where $Y_{ij}$ was the observed frequency (taxon count divided by library size) of the $j$th taxon in the $i$th sample, $\overline{Y}_j$ was the average frequency for the $j$th taxon across samples, $\delta_j$ was randomly drawn with value 1 or $-1$ with equal probabilities (and fixed across replicates of data), and $\mathcal{A}$ was the set of associated taxa. Note that the direction parameter $\delta_j$ had fixed value 1 for all $j$ in the simulations reported in [6], but was varied here to ensure (approximately) no association at taxa not selected into $\mathcal{A}$. In part, this represented our interest in testing individual taxa, which was not considered by [6]. Finally, we simulated the continuous trait as $X_i = \beta \text{scale}(C_i) + \epsilon_i$, where $\epsilon_i \sim N(0, 1)$ and scale(.) standardized the input vector to have mean 0 and standard deviation 1. This model tended to simulate weak associations for all associated taxa, although the abundant taxa generally had a higher impact.

Prior to analysis, we filtered out taxa that were present in fewer than 5 samples, which resulted in ∼460 taxa remaining in each simulated dataset. For evaluating sensitivity of detecting associated taxa, $\beta$ was set to 0.5 for S1–S5 in Model 1 and 5 for S1–S5 in Model 2. For evaluating power of the global test, $\beta$ was set to 0.2, 0.3, 0.3, 0.2, and 0.1 for S1–S5, respectively, in Model 1 and 0.5, 1, 5, 1, and 5 for S1–S5 in Model 2.

#### References

1. Hu YJ, Satten GA. Testing hypotheses about the microbiome using the linear decomposition model (LDM). Bioinformatics. 2020;p. bbtaa260, https://doi.org/10.1093/bioinformatics/btaa260.

2. Liu Y, Xie J. Cauchy combination test: a powerful test with analytic p-value calculation under arbitrary dependency structures. Journal of the American Statistical Association. 2020;115(529):393–402.

3. Wilson DJ. The harmonic mean p-value for combining dependent tests. Proceedings of the National Academy of Sciences. 2019;116(4):1195–1200.

4. Loughin TM. A systematic comparison of methods for combining p-values from independent tests. Computational statistics & data analysis. 2004;47(3):467–485.

5. Heard NA, Rubin-Delanchy P. Choosing between methods of combining p-values. Biometrika. 2018;105(1):239–246.

6. Zhao N, Chen J, Carroll IM, Ringel-Kulka T, Epstein MP, Zhou H, et al. Testing in microbiome-profiling studies with MiRKAT, the microbiome regression-based kernel association test. The American Journal of Human Genetics. 2015;96(5):797–807.

(a) Model 1

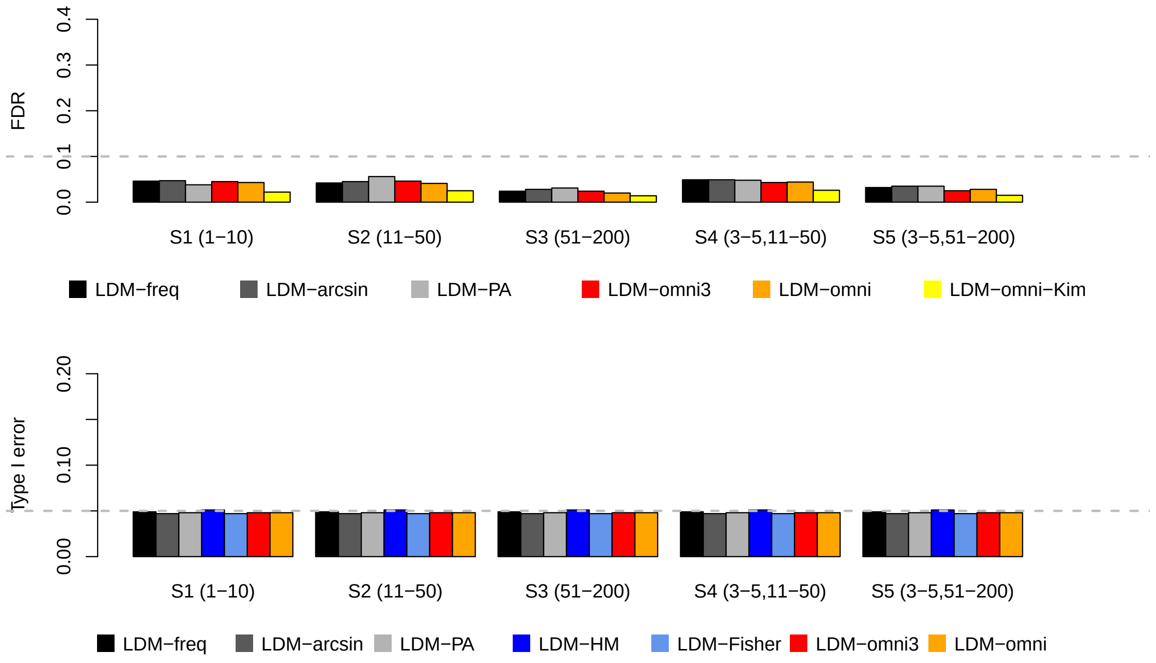


(b) Model 2

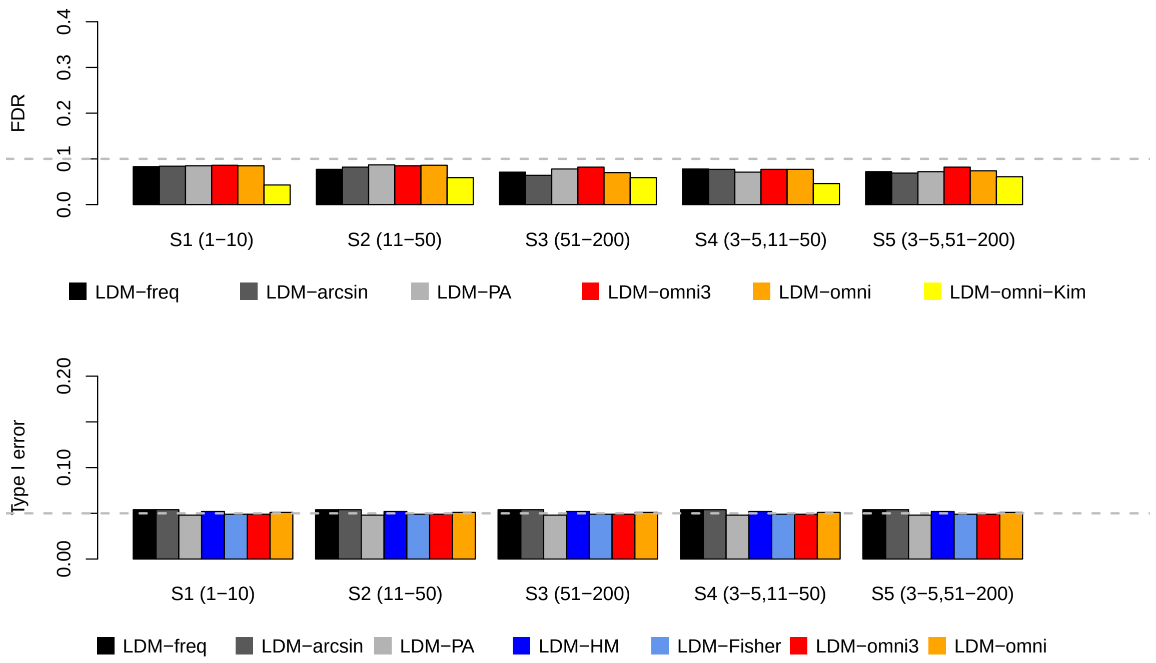


Figure S1: The gray dashed lines represent the nominal FDR 10% or the nominal type I error 0.05. The empirical FDR and type I error results were based on 1000 and 10000 replicates of data, respectively.

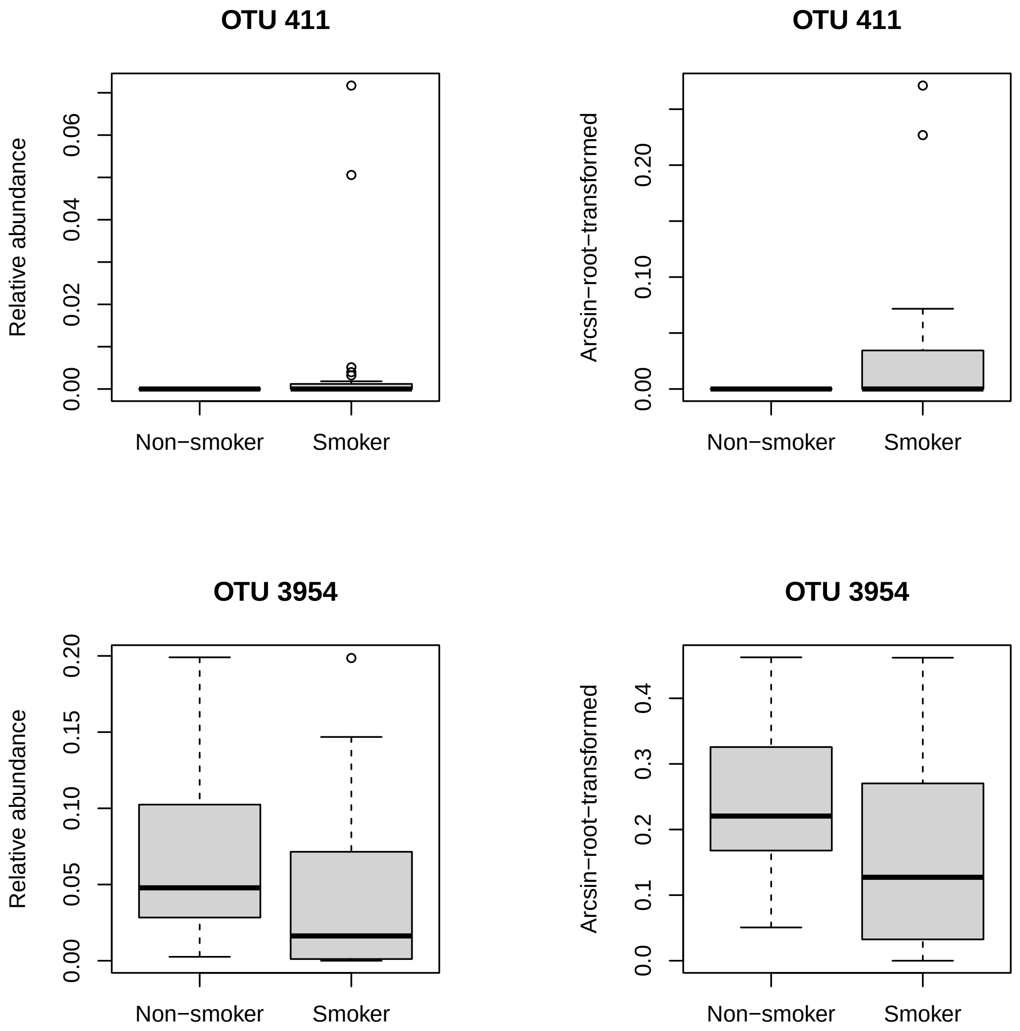


Figure S2: Distributions of the relative abundance data and the arcsin-root-transformed data of the two taxa, OTU 411 and OTU 3954, that were detected by LDM-omni3 but not by LDM-omni in analysis of the upper-respiratory-tract microbiome data.
